# Sequence Dependent Internucleosomal Interactions Dominate Array Assembly

**DOI:** 10.1101/2022.07.20.500866

**Authors:** Yaqing Wang, Tommy Stormberg, Mohtadin Hashemi, Anatoly B. Kolomeisky, Yuri L. Lyubchenko

**Author notes:** Corresponding author: Yuri L. Lyubchenko, Nebraska Medical Center, Omaha, NE 68198-6025. These authors contributed equally to this work (Yaqing Wang, Tommy Stormberg).

## Abstract

The organization of the nucleosome array is a critical component of the chromatin assembly into higher order structure as well as its function. Here we investigated the contribution of the DNA sequence and internucleosomal interactions to the organization of the nucleosomal arrays in compact structures using Atomic Force Microscopy. We assembled nucleosomes on DNA substrates allowing for the formation of tetranucleosomes. We found that nucleosomes are capable of forming constructs with the close positioning of nucleosomes with no discernible space between them, even in the case of assembled dinucleosomes. This morphology of the array is in contrast with that observed for arrays assembled with repeats of the nucleosome positioning motifs separated by uniform spacers. Simulated assembly of tetranucleosomes by random placement along the substrates revealed that nucleosome array compaction is promoted by the interaction of the nucleosomes. We developed a theoretical model to account for the role of DNA sequence and internucleosomal interactions in the formation of the nucleosome structures. These findings suggest that, in the chromatin assembly, the affinity of the nucleosomes to the DNA sequence and the strengths of the internucleosomal interactions are the two major factors defining the compactness of the chromatin.

## Introduction

DNA in eukaryotic cells is packaged into chromatin through extensive association with histone proteins (Roger D. Kornberg & Lorch, 1999; Van Holde & Zlatanova, 1995; Zhou et al., 2019). The nucleosome is the fundamental unit of chromatin, which regulates the readout and expression of the eukaryotic genome (Linking Chromatin Composition and Structural Dynamics at the Nucleosome Level, 2019; Roger D. Kornberg, 1974; Wilson & Costa, 2017). It is a DNA-protein complex with approximately 147 base pairs of DNA wrapped around a protein core complex known as the histone octamer (Elgin & Weintraub, 1975; R. D. Kornberg, 1977; McGhee & Felsenfeld, 1980). Canonical histone octamers consist of two copies of four core histone proteins, H2A, H2B, H3, and H4 (Isenberg, 1979). The positively charged histone octamers bind strongly to the negatively charged DNA. X-ray crystallography revealed the atomic structure of the nucleosome and explained how DNA is wrapped around histone octamers in a superhelix of approximately one and three-quarters turns (Luger et al., 1997).

The spatial organization of nucleosomes in chromatin continues to be a source of debate. Initially, it was proposed that nucleosomes condense into a 30-nm-diameter chromatin fiber supported by experiments with the use of electron microscopy (EM) or X-ray scattering analyses of chromatin extracted from various organisms (Chien & van Noort, 2009; Robinson et al., 2006; Schalch et al., 2005; Tremethick, 2007). Most recently, however, a study utilizing a combination of EM topography with a developed labeling method (ChromEMT) does not support the assembly of ordered 30-nm fibrils (Ou et al., 2017). Instead, they showed the assembly of 10-nm fibers in the cell that are not uniform; rather, they are heterogeneous and vary in diameter between 5 and 24 nm. Potential reasons for this discrepancy are discussed in the recent review article (Maeshima et al., 2019), in which the major role is given to electrostatics, as ionic strength for experiments *in vitro* and *in vivo* are very different. The authors also suggest that the absence of the 30-nm fiber formation can be due to nucleosome loss or irregular nucleosome spacing in native chromatin.

The findings in (Ou et al., 2017) are in line with publications (Luger & Hansen, 2005; Maeshima et al., 2010; Ohno et al., 2019) in which it has then been proposed that nucleosome fibers exist in a highly disordered, interdigitated state. What is the reason for such irregular spacing of nucleosomes? We have recently shown that the internucleosomal distance within dimers of nucleosomes varies depending on DNA sequences (Stormberg et al., 2019). No such effect is detected if repeats of such a highly sequence-specific DNA motif as the Widom 601 sequence are used in similar experiments (Filenko et al., 2012). Note that many structural studies of chromatin, including papers cited above, utilized repeats of the 601 motif (Boopathi et al., 2020).

Based on these studies we hypothesize that affinity to histone cores to DNA sequence can be a critical factor for the nucleosome array assembly stabilized by the internucleosomal interactions. To test this hypothesis, here we designed a DNA substrate containing a single instance of the specifically positioning 601 motif and an extended non-specifically positioning sequence from plasmid DNA or fully-non-specific DNA substrate. The experimental studies utilized single-molecule AFM, which can characterize the nanoscale structure of biological systems (Lyubchenko & Shlyakhtenko, 2016b; Miyagi et al., 2011; Pan et al., 2018; Stumme-Diers et al., 2019; Wang et al., 2020). We found that nucleosomes on such DNA templates form compact assemblies with close contacts between the nucleosomes. Monte Carlo simulation of random nucleosome placement along these substrates revealed that the compact assemblies observed experimentally occur at a much higher rate than expected with simple non-specific positioning. We propose a theoretical model according to which the affinity of the nucleosome core to the DNA sequence and the internucleosomal interactions are the two major factors defining the compact assembly of the nucleosome arrays.

## Results

The 601 DNA substrate consists of a positioning sequence, the 147 bp Widom 601 sequence, flanked by non-specific DNA sequences that can be wrapped by three or more histone octamers, as shown in Figure 1A. The Widom sequence provides a higher affinity for the histone octamers (Lowary & Widom, 1998), while the non-specific sequence provides an insight into a more biologically relevant aspect for the assembly of nucleosome array. The non-specific DNA substrate is identical in length to the 601 substrate, except that the 147 bp Widom 601 sequence is replaced by 147 bp of non-specific DNA, as shown in Figure 1B. The nucleosomes were assembled with the DNA substrate and histone octamer at a ratio of 1:5 DNA:octamer. A representative AFM image of the nucleosome assembly is shown in Figure 1C, where the bright features are the nucleosome cores, and the string-like strands are the unwrapped DNA in the complexes.

**Figure 1.**
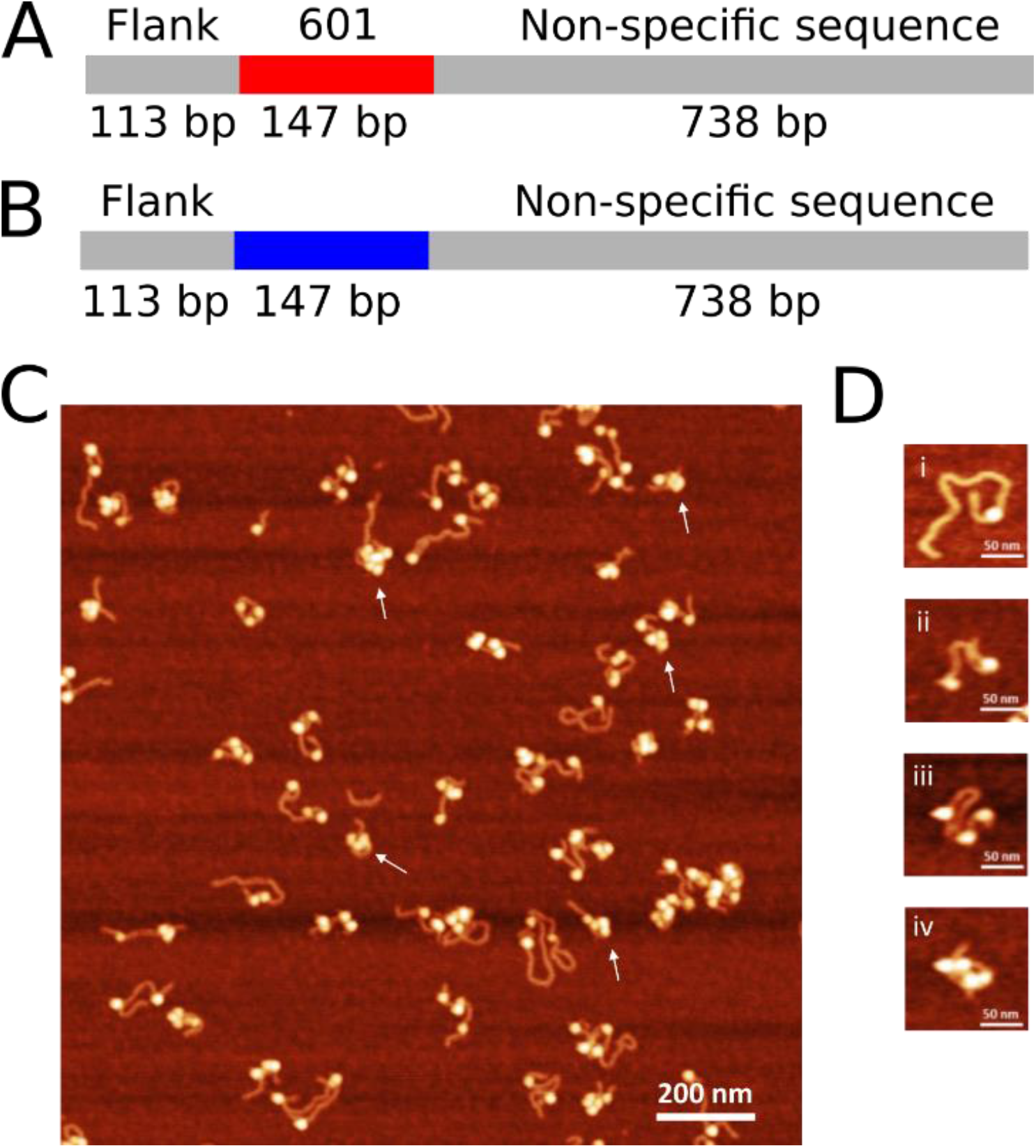
Nucleosome substrates and assembly. **A**. Schematic of 601 DNA substrate. **B**. Schematic of non-specific DNA substrate. **C**. Representative AFM image of nucleosomes assembled on the 601 DNA substrate. **D**. Selected images of different oligonucleosomes assembled on the 601 DNA substrate.

### Morphology of Nucleosome Arrays

From the acquired AFM images, we analyzed the morphology of the nucleosome arrays. Nucleosomes appear as bright globular features localized on DNA strands. The nucleosomes on the same DNA template can be separated, but immediately apparent was the compact morphology of nucleosome clusters. Examples of such compact arrangements of the nucleosomes are shown in Figure 1C, as indicated by the white arrows. These arrangements are irregular, resulting in nucleosome clusters of varying size and position. This result indicates that nucleosome arrays assembled on non-specific DNA adopt heterogeneous structure, as opposed to the ordered nucleosome arrays formed on substrates with repeated positioning DNA sequences imaged with AFM (Bash et al., 2001; Yodh et al., 1999, 2002).

Along with large nucleosome clusters, mono-, di-, tri-, and tetranucleosomes were observed. Representative snapshots of each type of nucleosome are shown in Figure 1D, with mononucleosome in frame (i), dinucleosome in frame (ii), trinucleosome in frame (iii), and tetranucleosome in frame (iv). We separated each type of nucleosome into groups based on the number of nucleosomes in the array and performed analysis. The yield of each oligonucleosome is shown in Table 1 (*n*=515). We found that mononucleosomes are the least commonly observed, while trinucleosomes and tetranucleosomes are the most common, indicating a preference for assembling higher order structures in our experimental setup.

**Table 1.** 601 Nucleosome Subpopulations

| | Yield ($n=515$) | Subpopulation | |
| --- | --- | --- | --- |
| **Free DNA** | 4.1% | N/A |  |
| **Mononucleosome** | 17.5% | N/A |  |
| **Dinucleosome** | 21.2% | 1-1 | 78.0% |
|  |  | 2 | 22.0% |
| **Trinucleosome** | 31.1% | 1-1-1 | 30.6% |
|  |  | 2-1 | 51.3% |
|  |  | 3 | 18.1% |
| **Tetranucleosome** | 26.2% | 1-1-1-1 | 6.7% |
|  |  | 2-1-1 | 18.5% |
|  |  | 2-2 | 13.3% |
|  |  | 3-1 | 41.5% |
|  |  | 4 | 20.0% |

We further segregated the groups of oligonucleosomes into subpopulations based on the proximity between nucleosomes on each array. Representative images of each subpopulation are shown in Figure 2. For dinucleosomes (Figure 2A), this resulted in two subpopulations; well separated (1-1) and compact (2) nucleosomes. For trinucleosomes (Figure 2B), three subpopulations exist; well separated (1-1-1), compact (3), and hybrid (2-1) nucleosomes. Tetranucleosomes (Figure 2C) contain five subpopulations; well separated (1-1-1-1), compact (4), and three hybrid structures, depending on the degree of compaction (2-1-1, 2-2, 3-1). The yields of the subpopulations are shown in Table 1. The well-separated dinucleosome accounts for 78.0% of all the dinucleosome population, while in the trinucleosome group, the well-separated subpopulation is observed in only 30.6% of cases. It decreases substantially in the tetranucleosome group, in which the well-separated tetranucleosomes represent only 6.7% of the population. The decreasing population of the well-separated subgroup in the higher-ordered structures further emphasizes that the compact morphologies are more favorable than the well-separated structures with distant spacing on this DNA substrate. Additionally, the “3-1” subpopulation counts for 41.5% of all tetranucleosomes observed, compared with 20.0% in the “4” subpopulation. This suggests that the trinucleosome may be the basic unit in the assembly of the nucleosomal array, rather than the tetranucleosome.

**Figure 2.**
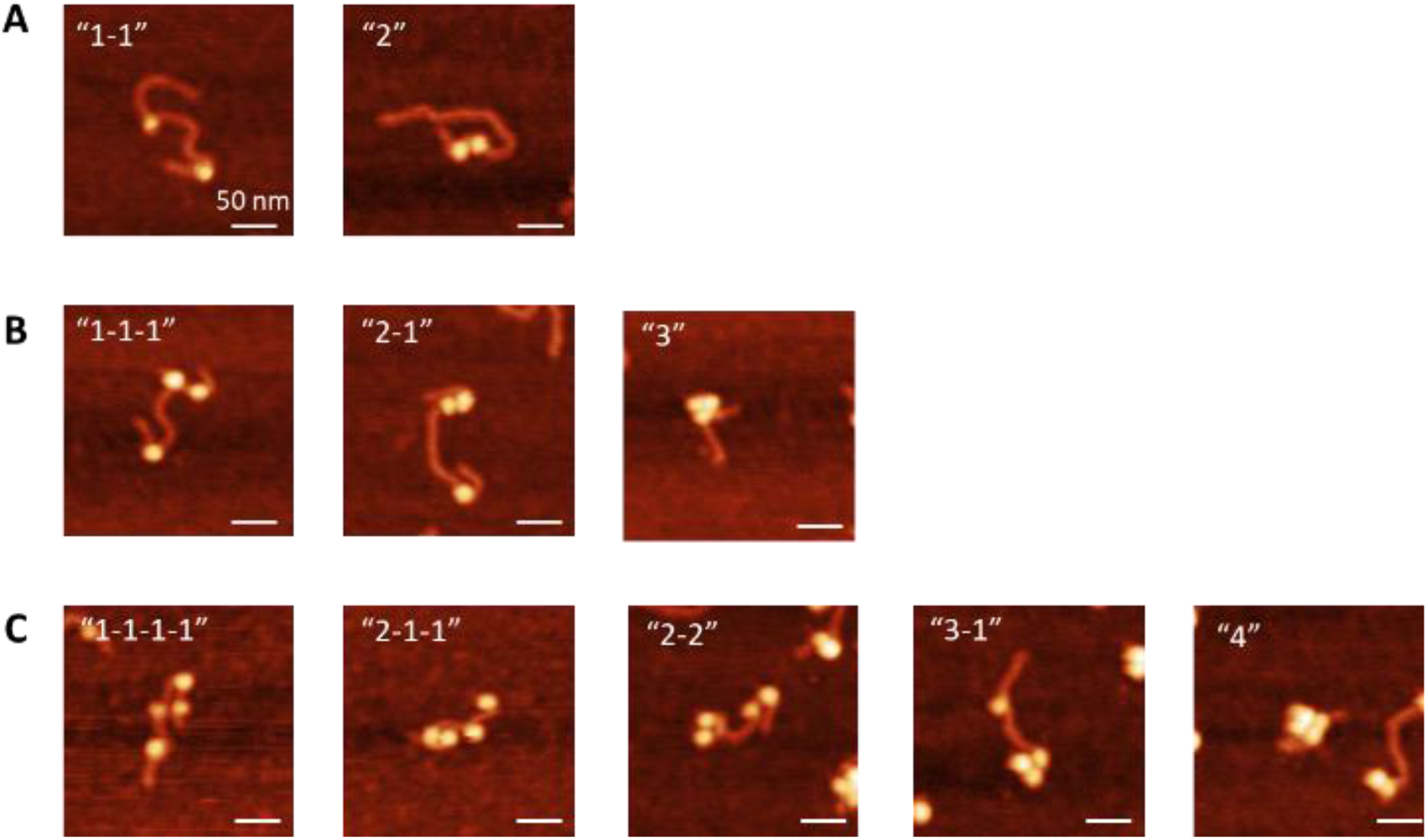
Subpopulations of oligonucleosomes. Scale bars indicate 50 nm.

### Internucleosomal Interaction within oligonucleosomes

In addition to the compact morphology, internucleosomal distances were also characterized. Figure 3 shows the data for both the well separated and compact dinucleosomes. The internucleosomal distances were measured from the center of one nucleosome to the center of the nearest neighboring nucleosome, as shown in Figure 3A,B. A representative image of measured distance for well separated nucleosomes (type 1-1) are shown in 3A, while an image of measured distance for compact nucleosomes (type 2) are shown in 3B. 10 nm was subtracted from each measured center-to-center distance to account for the size contributed by the histone core in each nucleosome. The results of these measurements were plotted and are shown in Figure 3C. We see that the separation between the well separated nucleosome and the compacted nucleosomes starts from 50 bp. This distance varies, with distances between nucleosomes as far as 650 bp apart, with no clear preference for positioning. In contrast, the distance of the compact nucleosomes is in the range of 0-40 bp, with a peak population centered at 28 ± 2 bp (SEM), as seen in Figure 3D. This indicates that nucleosomes within 30 bp of one another are most favorable for the internucleosomal interactions to compact the nucleosomes into proximity.

**Figure 3.**
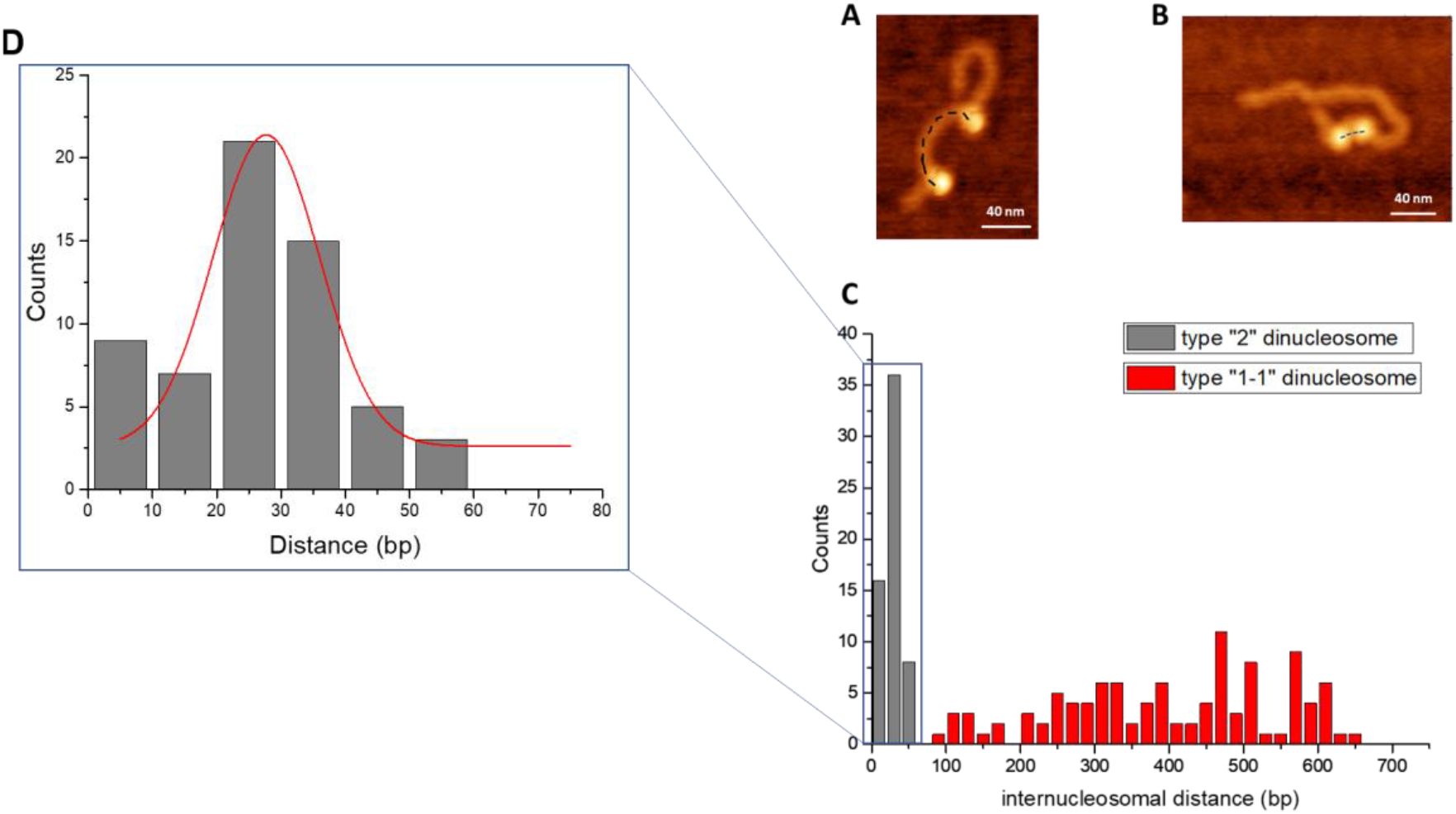
Analysis of internucleosomal distance in the dinucleosome population. **A**. Representative image of well separated dinucleosome (type “1-1”). **B**. Representative image of compact dinucleosome (type “2”). Scale bars indicate 40 nm. **C**. Histogram of internucleosomal distances, *n=*167. **D**. Distribution of internucleosomal distances in compact dinucleosomes indicate a peak value of 28 ± 2 bp (SEM).

The same measurements were performed on the “2-1” subpopulation of trinucleosomes, and the results can be seen in Figure S1. The distance between the two compacted nucleosomes is termed distance 1, while the distance between the well separated nucleosomes is termed distance 2. The combined data for internucleosomal distance, shown in S1B, indicates the same trend seen in the dinucleosome subpopulation. Compact nucleosomes are found in a very narrow region below 40 bp from one another, while well separated nucleosomes are found anywhere from 90 to more than 500 bp from one another. This result is in agreement with the results for dinucleosomes and further suggests that close proximity of nucleosomes is most favorable for the internucleosomal interactions to compact the nucleosomes into uniform structures.

### Nucleosomal Positioning

In order to determine the influence of the positioning 601 motif on the positioning and compaction of nucleosomes, we plotted the lengths of the free DNA flanks observed in our nucleosome arrays. The 601 motif was positioned 113 bp from one end of the DNA substrate; therefore, flank lengths of 113 bp would correspond to nucleosome positioning on the motif. The data for well separated dinucleosome flank length is shown in Figure 4. Interestingly, while a large portion of nucleosomes positioned at approximately 113 bp, nucleosome position varies, with nucleosomes positioning at or near the end of the DNA. This indicates that the preferential positioning of nucleosomes to the 601 motif is influenced by end binding and the total length of the DNA substrate.

**Figure 4.**
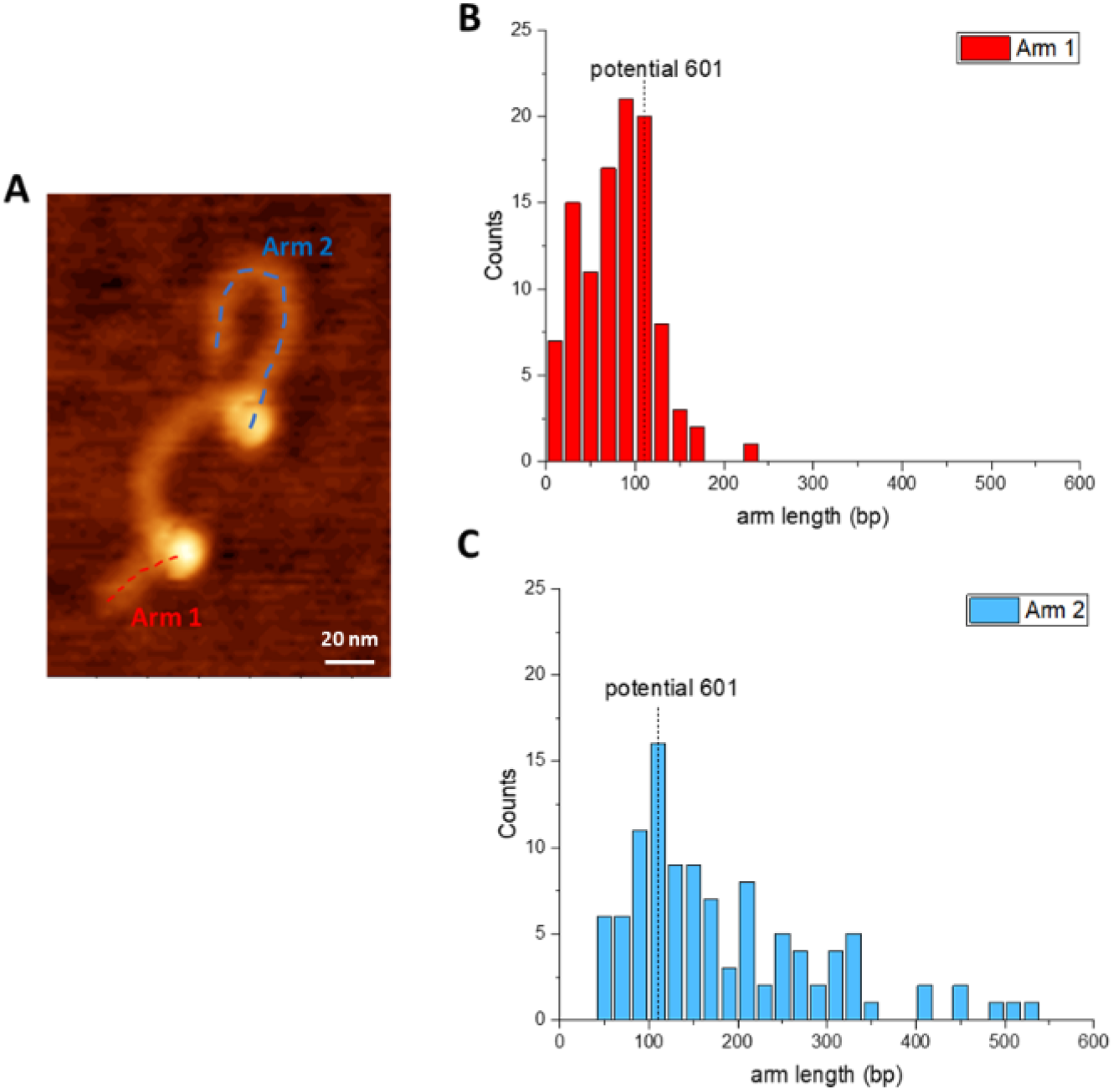
Analysis of flank DNA length in well separated dinucleosomes, *n*=106. Scale bar indicates 20 nm.

Compact structures of dinucleosomes and trinucleosomes both exhibited a similar positioning. The results are shown in Figure S2. The flank length of compact structures fall within the range of the 601 motif, but can vary as far as 250 bp away in the case of dinucleosomes. Compact trinucleosomes exhibit a lower range of positioning according to DNA flank length, but contain more structures positioned near the end of the DNA substrate. Overall, the presence of a single positioning nucleosome sequence does not appear to dictate the assembly pattern of other nucleosomes along the substrate, suggesting that array assembly is most dependent on interactions between nucleosomes.

### Nucleosome arrays with no positioning sequence

To further investigate the role of a positioning sequence in the assembly of higher order nucleosome structures, we prepared the DNA substrate comprising exclusively of non-specific DNA sequence of the same length as our substrate containing the 601 motif. The 147 bp 601 motif was replaced by 147 bp of non-specific DNA while maintaining the same sequence for the rest of the substrate. A schematic of the substrate is shown in Figure 1B, while a representative image of the assembled nucleosomes acquired through static AFM imaging is shown in Figure S3A. We found that, like nucleosomes assembled on the substrate containing the positioning sequence, the assembly was rather heterogeneous, including compact structures of varying sizes.

We again separated the oligonucleosomes into subpopulations; representative images of the oligonucleosome subgroups can be seen in Figure S3B-D. A table of the oligonucleosome yields is shown in Table 2 (*n*=498). It is notable that well separated subpopulation yields decrease in higher order structures, and the “3-1” population is the most common morphology-both observations match those of the substrate containing a single positioning sequence. However, on this substrate tri-and tetranucleosomes are more common compared with those assembled on the substrate containing a single positioning sequence (73.1% vs 60.3% of total complexes). Moreover, the overall population of well separated nucleosomes is diminished in the absence of a positioning sequence in observed dinucleosomes (65.5% vs 86.3%) and in tetranucleosomes (6.8% vs 10.7%). This result indicates that the absence of a positioning sequence may allow nucleosomes to move more freely along the substrate, resulting in more internucleosomal interactions and therefore more compact structures.

**Table 2.** Non-specific Nucleosome Subpopulations

|  | Yield (*n*=498) | Subpopulation |  |
| --- | --- | --- | --- |
| **Free DNA** | 4.0% | N/A |  |
| **Mononucleosome** | 5.4% | N/A |  |
| **Dinucleosome** | 17.5% | 1-1 | 65.5% |
|  |  | 2 | 34.5% |
| **Trinucleosome** | 46.4% | 1-1-1 | 34.6% |
|  |  | 2-1 | 47.2% |
|  |  | 3 | 18.2% |
| **Tetranucleosome** | 26.7% | 1-1-1-1 | 6.8% |
|  |  | 2-1-1 | 28.6% |
|  |  | 2-2 | 12.0% |
|  |  | 3-1 | 30.8% |
|  |  | 4 | 21.8% |

### Simulations of nucleosome positioning

We simulated the random placement of nucleosomes along the substrates to assess the impact of sequence and internucleosomal interactions in nucleosome proximity compared with random placement. For the non-specific sequence, tetranucleosomes were simulated by randomly placing four particles occupying 147 bp along the substrate with no allowance for overlap. For the 601 sequence, we simulated the substrate with the region containing the 601 motif (from 113-259 bp) as occupied to mimic a homogenously bound nucleosome in that region; three particles were then randomly placed along the remainder of the substrate, reflecting a randomly assembled tetranucleosome array with one nucleosome bound to the 601 region. We also simulated a shifted version of the 601 substrate - the region from 148-294 bp was pre-occupied, while three particles were randomly placed along the rest of the substrate. This setup allowed for a nucleosome to bind upstream of the excluded region. The results were analyzed and grouped into subpopulations based on their proximity to the next nucleosome along the array.

Nucleosome pairs within 30 bp of one another were considered compact, while those greater than 30 bp from one another were considered well separated, as determined from the data shown in Figure 3. The yield of subpopulations is shown in Table 3 (N=1000). Non-specific denotes the non-specific DNA substrate simulation, 601 denotes the 601 substrate simulation, and 601 shifted denotes the shifted 601 substrate simulation. There are stark differences between the experimental and simulation data.

**Table 3.** Simulated Tetranucleosome Subpopulations

| Subpopulation | Nonspecific Yield | 601 Yield | 601 Shifted Yield |
| --- | --- | --- | --- |
| **1-1-1-1** | 33.2% | 29.7% | 24.8% |
| **2-1-1** | 51.1% | 50.6% | 52.4% |
| **2-2** | 4.5% | 4.6% | 5.7% |
| **3-1** | 10.3% | 14.8% | 16.3% |
| **4** | 0.9% | 0.3% | 0.8% |

Simulated data reveals the “2-1-1” subpopulation to be the most common morphology along all three setups, representing approximately half the dataset in all three cases. The “4” subpopulation of simulated tetranucleosomes is particularly low, representing less than 1% of the dataset in all three cases, dropping as low as 0.3% for 601 tetranucleosomes. Yet, experimental data shows that the “2-1-1” subpopulation represents as much as 28.6% of the dataset for non-specific tetranucleosomes. The “4” subpopulation represents slightly less; 21.8% for non-specific tetranucleosomes. Likewise, for 601 tetranucleosomes, experimental data reveals a mere 17.5% of tetranucleosomes in the “2-1-1” conformation, and 19.4% are observed in the “4” conformation. A full comparison of all three simulations compared with the experimental datasets can be seen in Tables S1 and S2. Overall, the simulation data show a significant preference for well separated nucleosomes compared to the experimental data. In fact, the assembly of the “4” conformation is more than 24 times more likely in experiments (with either substrate) compared with their simulated counterparts, with the “4” conformation in 601 tetranucleosomes being nearly 65 times more likely (Tables S1 and S2). This indicates that nucleosome positioning along the DNA substrate is not random - it is very likely influenced by internucleosomal interactions.

### Theoretical model for nucleosome interaction and positioning

To understand the dynamics of interactions between nucleosomes and DNA as well as internucleosomal interactions, we develop a simple chemical model that allows us to extract the most relevant properties of these complex processes. Based on the experimental constructs, we consider two DNA molecules of length *L*=998 bp, with and without a single 601 segment. Nucleosomes can bind to the DNA with an effective energy *E*_*ns*_ (in units of *k*_*B*_*T*), if the association happens to the non-specific region, or with an effective energy *E*_*601*_, if the association happens to the 601 sequence. Each association covers *l*=147 bp of the DNA length. In addition, two DNA-bound nucleosomes that are located at the distances less than *d*=30 bp are assumed to have internucleosomal interactions *E*_*int*._ This cutoff distance is chosen based on the experimental data for dinucleosomes in Figure 3 that show the peak in the distribution of internucleosomal distances as being 28 ± 2 bp (SEM).

Experimental data showed that there are a total of 12 observable chemical states in the system: one state when DNA is totally free, one state with a single bound nucleosome, two states with two bound nucleosomes (with and without internucleosomal interaction); three states with three bound nucleosomes; and five states with four bound nucleosomes. Because of long times of observations during AFM experiments, it is reasonable to assume that the system reaches an overall chemical equilibrium. This allows us to significantly simplify the analysis to only few states and transitions between them.

Following this strategy, we first consider the transitions between the free DNA (state 0) and mononucleosome (state 1) for non-specific DNA. We define the equilibrium probabilities of these states as *P*_*0*_ and *P*_*1*_, respectively. The condition of chemical equilibrium then requires that

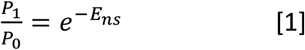

Using the data from Table 2, one can easily obtain that the effective free energy difference is *E*_*ns*_*=0.3 k*_*B*_*T*. Extrapolating based on this, one can obtain the energy of the mononucleosome state, 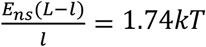. This means that interactions between nucleosomes and non-specific segments of DNA are relatively weak, allowing nucleosomes to be quite mobile.

The equilibrium between free DNA and mononucleosomes for the system with the 601 sequence is more involved because the nucleosome can associate to this 601 sequence (we call it a state 1*) or it can associate to any other non-specific sequence (we call it a state 1). The probabilities, at chemical equilibrium, between these states can be written as

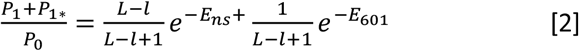

This equation reflects the fact that only one site of available *L-l+1* binding sites is a special one, and 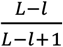 of them are non-specific bindings. Using the already obtained value for *E*_*ns*_*=0.3 k*_*B*_*T* and the parameters for *L* and *l*, we obtain that *E*_*601*_*=8.3 k*_*B*_*T*. This shows that the interactions between nucleosomes and the special 601 positioning sequence are very strong, preventing the mobility of the bound nucleosome in this case. Furthermore, the estimate is in good agreement with experimental results published by other groups (Van Der Heijden et al., 2012).

We then explore the inter-particle interactions for bound nucleosomes. We consider the equilibrium distributions of two different states for dinucleosomes on non-specific DNA. To simplify the analysis, let us neglect the end effect of the finite length of the DNA molecule. Then, if one nucleosome is already bound, there are *L-3l+2* sites where the second nucleosome might bind. But *2d* of them will exhibit an additional internucleosomal interaction energy. This leads to the fraction of associations that lead to the state 2 (two bound, interacting nucleosomes) to be

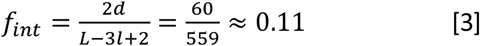

Then the fraction of binding events that lead to the state 1-1 (two bound, non-interacting nucleosomes) is *1-f*_*int*_*=0.89*. The equilibrium distribution of states for dinucleosomes then gives rise to the following

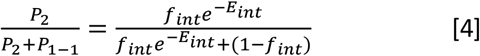

Using data from Table 2, we estimate that *E*_*int*_ *∼ 1.5 k*_*B*_*T*, or 0.9 kcal/mole, which is very close to internucleosomal interaction energy obtained in other experimental studies (Widlund et al., 2000).

The robustness of our analysis can be tested by applying it to evaluate the fraction of states for the dinucleosomes in the case of the special 601 positioning sequence, that was measured independently in our experiments. Because of strong interactions with the 601 sequence, it is reasonable to assume that one nucleosome is always bound to this special site. The second nucleosome can associate to one of *L-2l-l*_*0*_*+1=592* sites. *l*_*0*_*=113* bp is the distance from the 601 segment to one end of DNA, which also imposes that the second nucleosome can only bind to one side of the already bound nucleosome. This gives the following fraction of binding sites that lead to state 2 (two interacting nucleosomes)

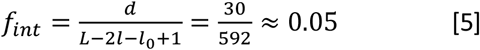

Substituting this result into Eq. [4] predicts that ∼18.8% of dinucleosomes formed on the substrate with the special 601 positioning sequence, would correspond to the state 2, which is comparable to experimentally measured value of 21.9% for state 2.

## Discussion

We used AFM to characterize the role of DNA sequence and internucleosomal interactions on the assembly of the nucleosome array. We assembled oligonucleosomes on two DNA substrates; the first comprising 998 bp of DNA with a single nucleosome positioning motif located near the end of the DNA (601 DNA), and the second of the same length, but with no regions of high nucleosome specificity (non-specific DNA), which mimics natural DNA. The most striking feature of the AFM images of the arrays assembled on these DNA templates is the ability of nucleosomes to assemble in clusters with close proximity of neighboring nucleosomes. Such clusters with two, three and four nucleosomes comprise more than 90% of tetranucleosome arrays, whereas species with distant locations of nucleosomes comprise only 6.7% on 601 DNA (Table 1) and 6.8% on non-specific DNA (Table 2). A similar effect of the nucleosome clustering is observed for the trinucleosome assembles, although a well-separated 1-1-1 arrangement for trinucleosome is more likely, 30.6% and 34.6% for 601 DNA and non-specific DNA, respectively. This finding is in a sharp contrast with AFM studies of arrays assembled on repeats of nucleosome-positioning sequences in which arrays of well-separated nucleosomes are the predominant features (Bash et al., 2001; Bussiek et al., 2007; Yodh et al., 1999, 2002).

To quantitatively characterize the internucleosomal interactions, we measured internucleosomal distances for dinucleosome and 2-1 trinucleosome assemblies, for which this parameter can be measured using AFM images. Quantitative analysis for both assemblies shown in Figs. 3 and S1 reveal a number of features of the nucleosome assemblies. These data graphically demonstrate that nucleosomes can be tightly clustered in dinucleosome assemblies or separated from each other in a broad range. Importantly, the nucleosomes in the dinucleosome populations, separated at a peak distance as small as 30 bp, showed compact assemblies in which 24% of species display an internucleosomal distance of less than 10 bp. Overall, the distribution is narrow, with a standard deviation of 13 bp. Monte Carlo simulations failed to identify such frequent and close contacts when nucleosomes were allowed to position freely along the substrates (Tables S1, S2). These data point directly to the role of internucleosomal interactions. According to numerous publications (paper (Lyubartsev et al., 2015) and references therein), histone tails play a critical role in the formation of tight internucleosomal contacts, as post-translational modifications such as acetylation can decrease the nucleosome clustering, including the assembly of dinucleosomes studied with AFM (Bash et al., 2001).

According to the molecular modeling of nucleosome arrays (Lyubartsev et al., 2015), the density of the nucleosome packing depends on the orientation of the nucleosomes in which the tightest distance (less than 1nm between the nucleosomes) corresponds to stacking of the disk-type particles. The distance can be larger for other orientations of nucleosomes, and the variation of the entry-exit angles between the adjacent nucleosomes, defined by the size of linker DNA between the nucleosome, contributes to the orientation of the nucleosome and hence the internucleosomal distance. We assume that the orientation factor can explain the breadth of the internucleosomal distance distribution in our experiments. Note however, that histone tails not only bridge the nucleosomes, but also can hinder the formation of the tight internucleosomal contacts. We have recently shown that truncation of histone H4 can shift the internucleosomal distance for the dinucleosome constructs to smaller distances (Stormberg, Vemulapalli, et al., 2021).

Clustered dinucleosomes were visualized before (Bussiek et al., 2007; Yodh et al., 1999, 2002), although these were minor species due to the use of DNA templates with positioning sequence repeats, so the nucleosomes primarily occupied the nucleosome-specific motifs. The assembly of dinucleosomes in(Yodh et al., 1999, 2002) was explained by a relatively low energetic preference for assembly of nucleosomes on favored motifs compared with non-favored ones, however, three-nucleosome clusters have not been observed in those papers, nor four-nucleosome clusters, which in our case were observed with high yield. Our theoretical model and estimates made with the comparison with experiments provide the following explanation for the nucleosome clustering. The affinity of nucleosome to non-specific DNA sequence is as low as 0.3 k_B_T. Given that the internucleosomal attraction is relatively high, *E*_*int*_ *∼ 1.5 k*_*B*_*T*, or 0.9 kcal/mole, as per our calculations above and previous publications (Widlund et al., 2000), the assembly of clusters is a favorable process. However, affinity of nucleosome to positioning sequences can prevent the assembly of nucleosome clusters. Importantly, our theoretical analysis explains that differences in distributions of states in dinucleosomes are due to different interaction strengths between the nucleosomes and non-specific and 601 positioning DNA segments. Thus, the morphology of nucleosome array is defined by the DNA sequence, so one can expect clusters along with non-clustered segments in the array. However, natural nucleosomal DNA contains only modest sequence periodicity (Shrader & Crothers, 1989; Yodh et al., 2002), so clustering of nucleosomes should be the most representative morphology of the chromatin. AFM images of trinucleosome and tetranucleosome clusters did not reveal geometric features of periodic solenoid model of chromatin.

The non-regular morphology of tri-and tetra-nucleosome arrays observed in this study better fit the model of irregular chromatin structure (Ou et al., 2017). Importantly, the assembly of nucleosome in clusters can be an important factor in the gene regulation. Promoter regions in eukaryotes do not show a high affinity to nucleosome assembly (Lee et al., 2007; Mavrich et al., 2008; Ozsolak et al., 2007) allowing for the nucleosome clustering. Such clusters would create a hurdle for transcription factors to bind regulatory regions as well as for RNA polymerase to transcribe genes.

## Materials and Methods

### DNA Substrate

The 601 DNA substrate used in nucleosome assembly contains the 147 bp Widom 601 sequence flanked by plasmid DNA, 113 bp and 738 bp in length (shown in Figure 1A). It is generated from PCR using a plasmid vector pUC57 with the forward primer (5’-GATGTGCTGCAAGGCGATTAAG-3’) and the reverse primer (5’-GGGTTTCGCCACCTCTGAC-3’). The 998 bp non-specific DNA substrate used is the same length as the 601 substrate, except that the 147 bp Widom 601 sequence is replaced by 147 bp of non-specific DNA (shown in Figure 1B). The DNA substrates were concentrated from the PCR product and purified using gel electrophoresis. DNA concentration was then determined using NanoDrop Spectrophotometer (ND-1000, Thermo Fischer) before being used for nucleosome assembly.

### Nucleosome Assembly

Nucleosomes were assembled using a gradient dilution method optimized from our previous research (Stormberg et al., 2019; Stormberg, Filliaux, et al., 2021; Stumme-Diers et al., 2018, 2019). Recombinant human histone octamers were purchased from The Histone Source (Fort Collins, CO) for assembly. Before assembly, histones were dialyzed against the initial dialysis buffer (10 mM Tris pH 7.5, 2 M NaCl, 1 mM EDTA, 2 mM DTT) at 4°C for 1 hour. DNA (25 pmol) was then mixed with the histone octamer at a molar ratio of 1:5. The total volume of the mixture was adjusted to 10 μL with 5 M NaCl and DDI H_2_O so that the starting concentration of NaCl in the reaction is 2 M. The mixture was diluted with dilution buffer (10 mM Tris pH 7.5) using a syringe pump (0.07 μL/min for 1000 min) to gradually decrease the salt concentration to 0.25 M NaCl, allowing the histone to bind the DNA and form the nucleosome core particle. The nucleosomes were then dialyzed for 1 hour against a fresh low salt buffer (10 mM Tris pH 7.5, 2.5 mM NaCl, 1 mM EDTA, 2 mM DTT) before being diluted to 300 nM and stored at 4°C. The final concentration of the nucleosome was adjusted to 2 nM right before deposition using imaging buffer (10 mM HEPES pH 7.5, 4 mM MgCl_2_).

### AFM Imaging and Data Analysis

Sample preparation for AFM imaging was performed as described in (Lyubchenko & Shlyakhtenko, 2016a; Stormberg et al., 2019; Stumme-Diers et al., 2019). The nucleosome sample was deposited onto the APS-functionalized mica and incubated for 2 minutes at room temperature. After that, the sample was rinsed with DDI H_2_O and dried with a gentle argon flow. The samples were stored in vacuum before being imaged.

Images were acquired using tapping mode under ambient conditions on a MultiMode 8, Nanoscope V system (Bruker, Santa Barbara, CA) using TESPA probes (320 kHz nominal frequency and a 42 N/m spring constant) from the same vendor. The dry sample AFM images were analyzed using the FemtoScan Online software package (Advanced Technologies Center, Moscow, Russia).

DNA contour length analysis was performed by tracing the DNA from one free end to the other. The mean measured contour length of free DNA on an image was divided by the known length of the given substrate, yielding the conversion factor. Flank measurements for the nucleosomes were obtained by measuring from the DNA end to the center of the nucleosome for both arms. 5 nm was subtracted from each measured flank length to account for the contribution by the histone core. The flank length measurements were divided by the calculated conversion factor to convert from nm to base pair values.

The distinction between compact and well separated nucleosome subpopulations was performed visually by separately grouping nucleosomes that were observed to be in direct contact from those with clearly distinguishable DNA between nucleosome core complexes. Internucleosomal distance cutoff used in simulations and theoretical calculations was determined by measuring the center-to-center distance of dinucleosomes, subtracting 10 nm to account for the contribution of each histone core, and dividing by the calculated conversion factor to convert from nm to bp. The histograms were generated using OriginPro software (OriginLab Corporation, Northampton, MA, USA).

### Monte Carlo Simulations and Data Analysis

Monte Carlo simulations for random placement of 147 bp segments on a 998 bp DNA substrate were performed using bedtools (v2.30.0) (Quinlan & Hall, 2010). Each simulation randomly placed four segments on the DNA substrate, prohibiting overlap of the segments; simulations were repeated 1,000 times for each substrate. Simulations were performed for substrates with no sites occupied, one site occupied (113-259 bp) corresponding to the substrate with positioning sequence, and a shifted occupied site (148-294 bp) corresponding to a substrate with positioning sequence allowing nucleosome assembly on both side of the specific sequence. Data was compiled in Excel and grouped into subpopulations based on their proximity to the next nucleosome along the array using the cutoff above. Nucleosome pairs within 30 bp of one another were considered compact, while those greater than 30 bp from one another were considered well separated.

## ACKNOWLEDGMENTS

The authors thank the Lyubchenko lab members for useful insights in the data interpretation.

## SUPPORTING INFORMATION

**Figure S1.**
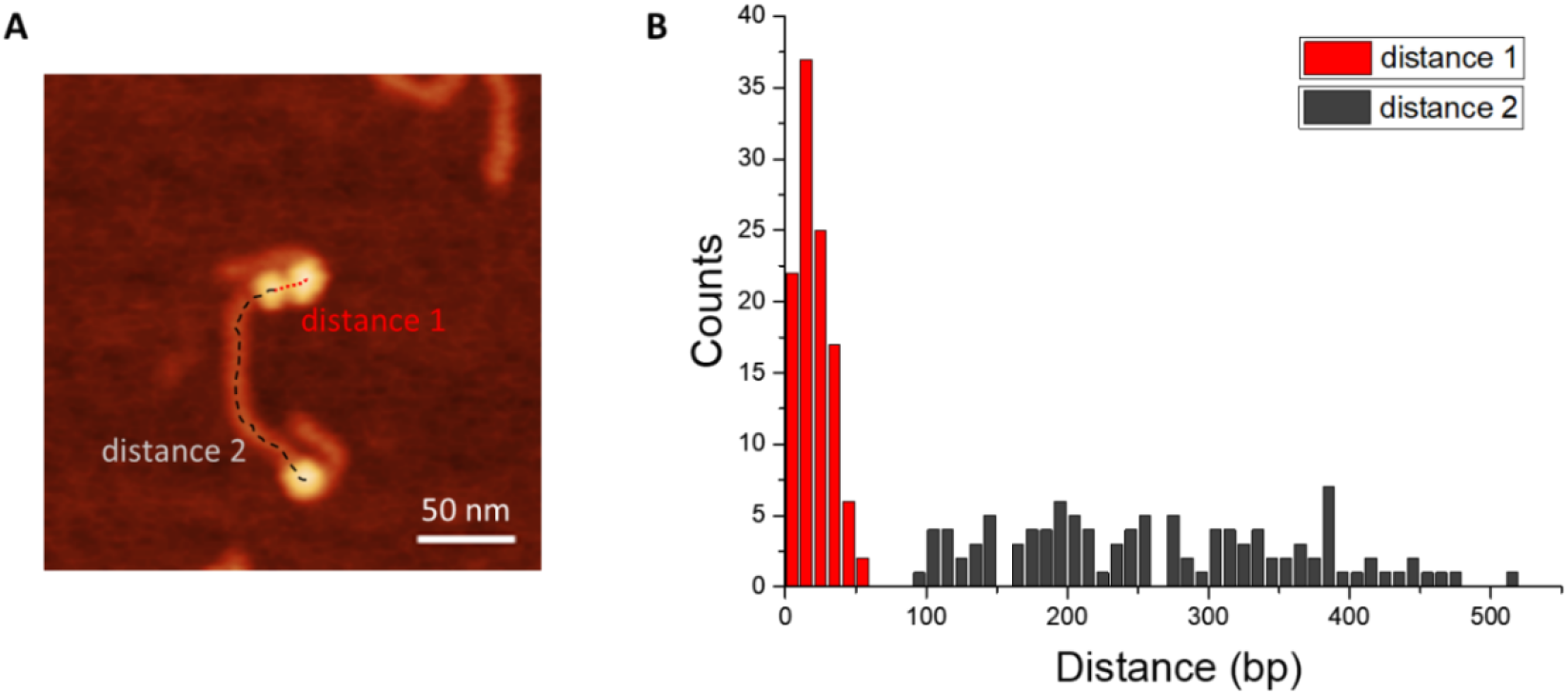
Analysis of the internucleosomal distance in the 2-1 trinucleosome population, *n*=109. Scale bar indicates 50 nm.

**Figure S2.**
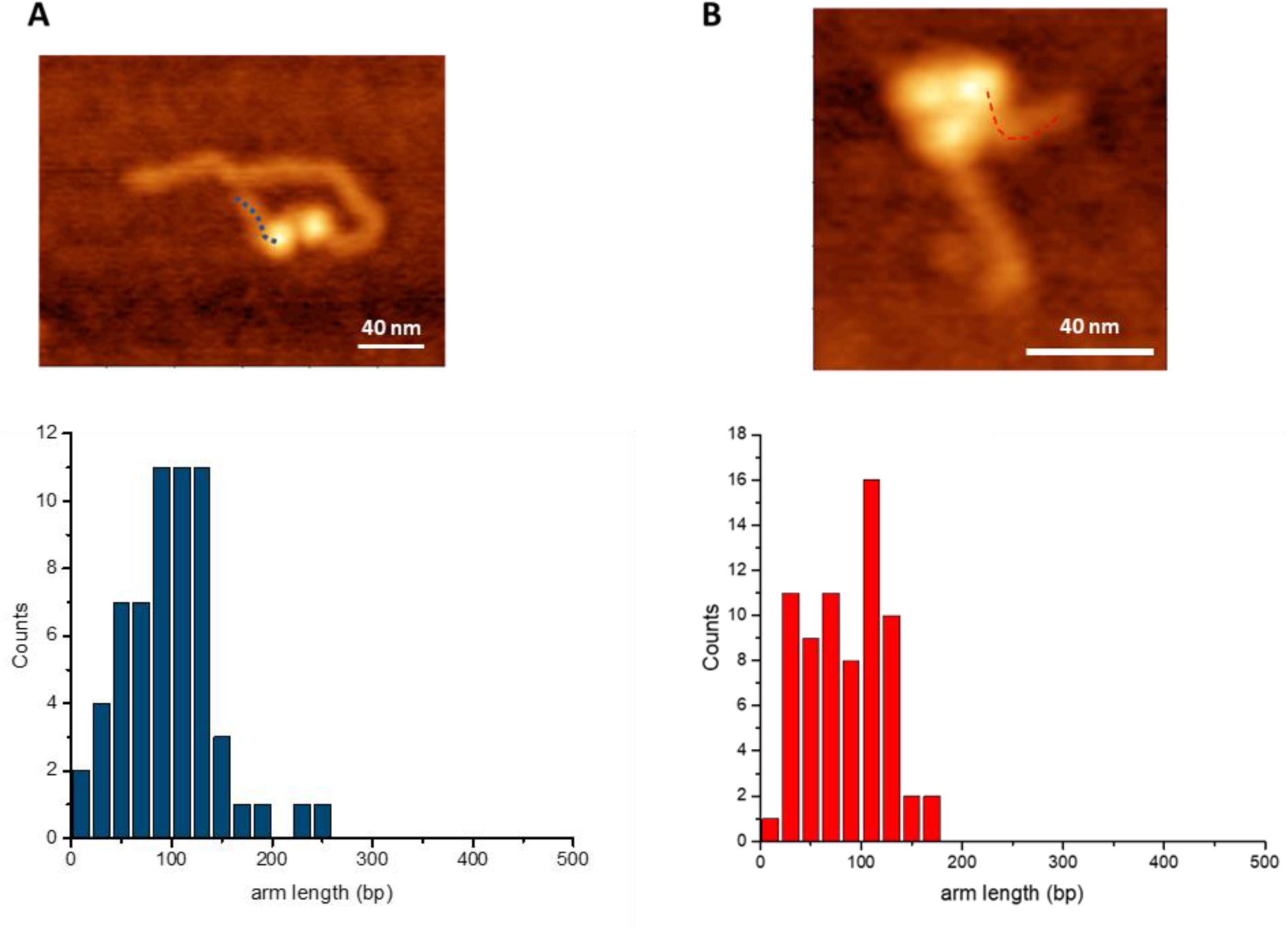
Analysis of flank DNA length in compact dinucleosomes, *n*=60 **(A)** and trinucleosomes, *n*=70 **(B)**. Scale bars indicate 40 nm.

**Figure S3.**
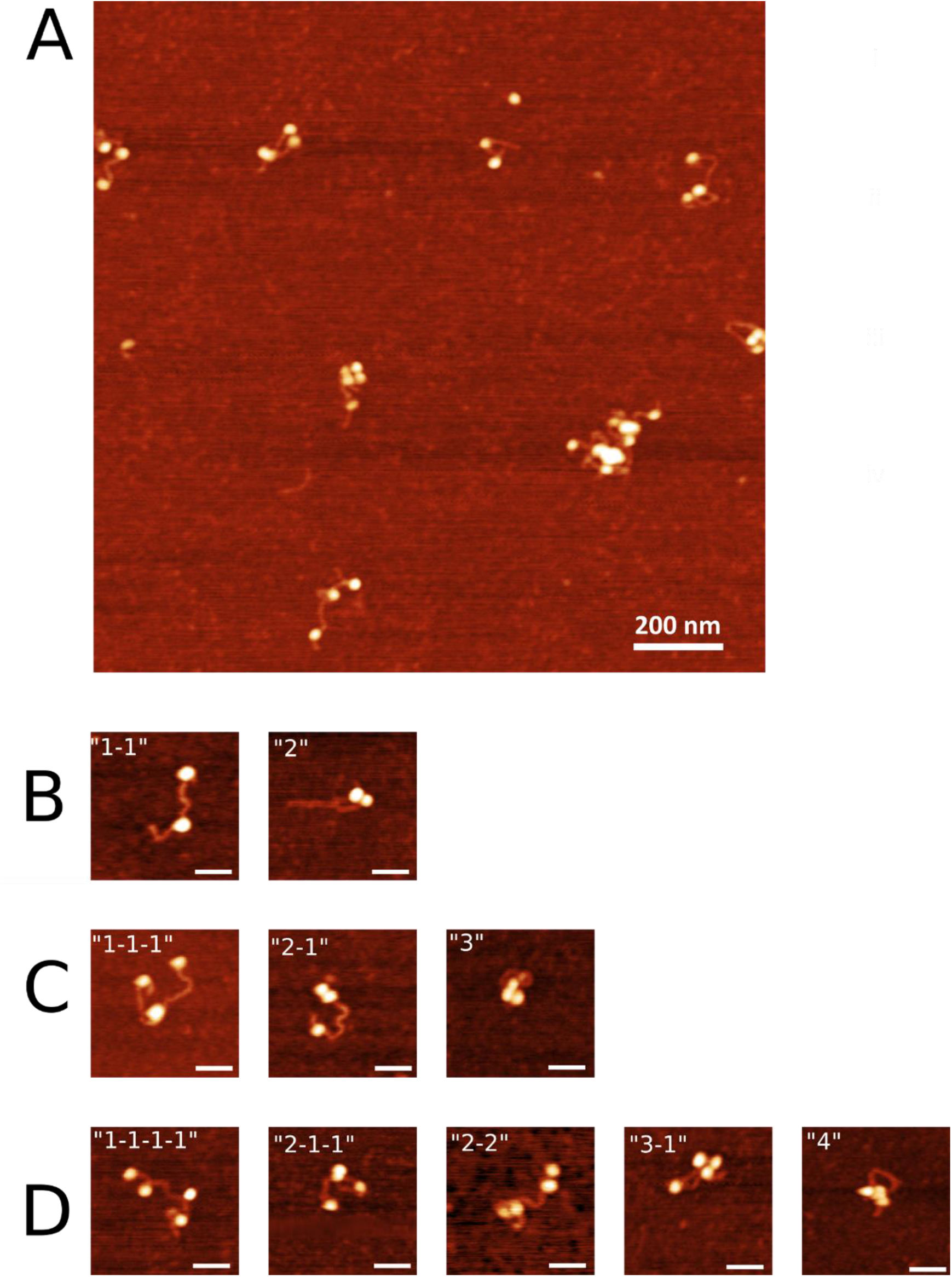
Nucleosomes assembled on non-specific DNA. **A**. Representative AFM image. **B-D**. Subpopulations of oligonucleosomes. Scale bars indicate 50 nm.

**Table S1.**
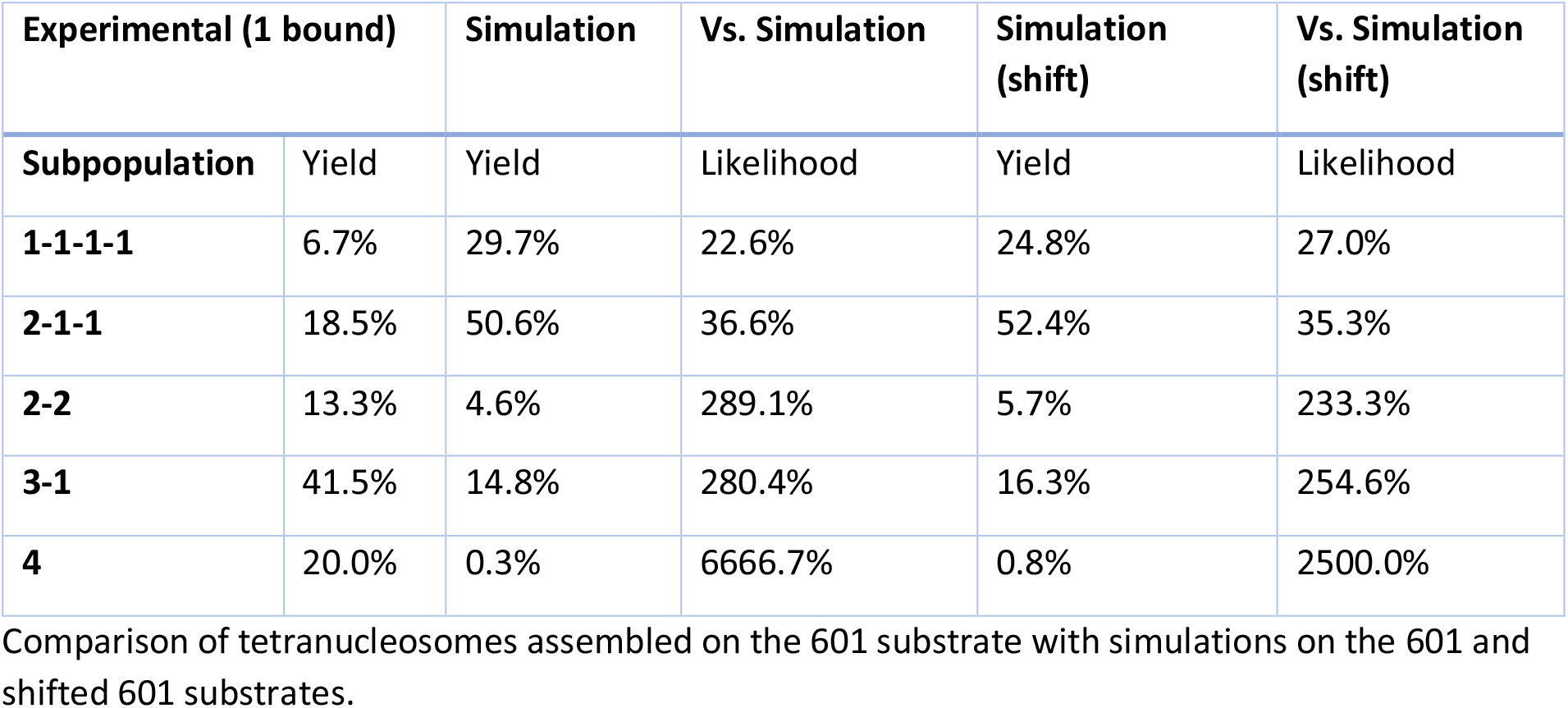
601 Tetranucleosome Comparison

| Experimental (1 bound) |  | Simulation | Vs. Simulation | Simulation (shift) | Vs. Simulation (shift) |
| --- | --- | --- | --- | --- | --- |
| Subpopulation | Yield | Yield | Likelihood | Yield | Likelihood |
| 1-1-1-1 | 6.7% | 29.7% | 22.6% | 24.8% | 27.0% |
| 2-1-1 | 18.5% | 50.6% | 36.6% | 52.4% | 35.3% |
| 2-2 | 13.3% | 4.6% | 289.1% | 5.7% | 233.3% |
| 3-1 | 41.5% | 14.8% | 280.4% | 16.3% | 254.6% |
| 4 | 20.0% | 0.3% | 6666.7% | 0.8% | 2500.0% |

Comparison of tetranucleosomes assembled on the 601 substrate with simulations on the 601 and shifted 601 substrates.

**Table S2.**
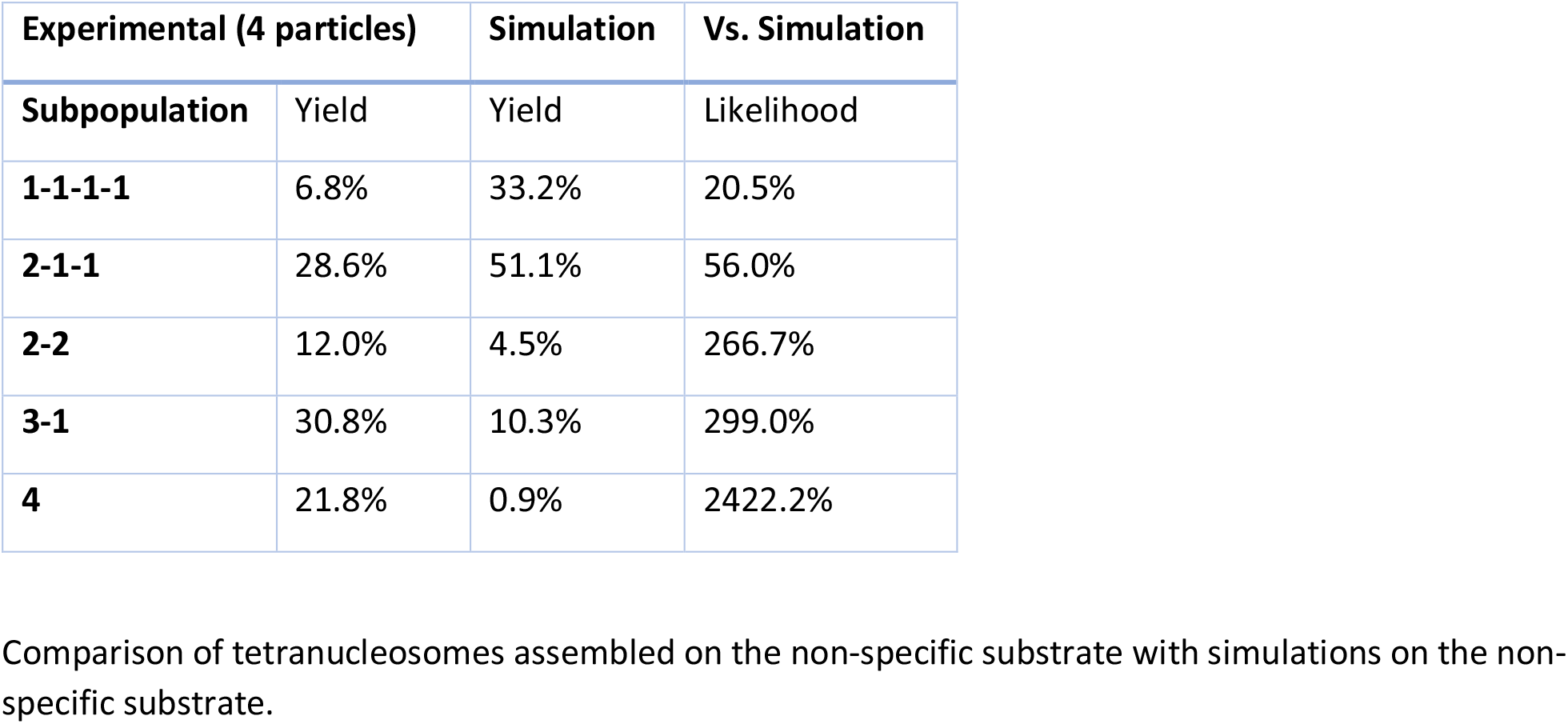
Non-specific Tetranucleosome Comparison

